# deCYPher: Star Allele-Resolution Computational Framework of Pharmacogenes for Haplotype-Resolved Long-Read Assemblies

**DOI:** 10.1101/2025.10.13.681303

**Authors:** Ting-Yu Chang, Yu-Shi Liu, Hong-Sheng Lai, Tsung-Kai Hung, Hsin-Fu Lin, Yu-Hui Lin, Chia-Lang Hsu, Ya-Chien Yang, Chein-Yu Chen, Pei-Lung Chen, Jacob Shu-Jui Hsu

## Abstract

Although existing next-generation sequencing (NGS) tools, such as Aldy and Cyrius, have been applied for allele typing, they cannot achieve complete accuracy due to various genomic challenges including pseudogenes, structural variations, hybrid genes, copy number variations, and gene deletions. These complexities make accurate pharmacogene interpretation more challenging, despite the crucial role pharmacogenomics plays in precision medicine. We developed **deCYPher**, a tool that generates personalized pharmacogenomic reports from haplotype-resolved assemblies. The tool enables analysis of all PharmVar 1A level genes, such as *CYP2B6, CYP2C9, CYP2C19, CYP2D6, CYP3A5, CYP4F2, DPYD, NUDT15*, and *SLCO1B1*. Applied to all HPRC haplotypes (including both release 1 and release 2 data), deCYPher demonstrated high accuracy in resolving complex gene structures. In the case of CYP2D6, release 1 identified 6% gene multiplications, 6% full gene deletions, and 4% CYP2D6/CYP2D7 hybrids. By contrast, release 2 demonstrated an increased prevalence of multiplications (14%) and hybrids (11%), while the frequency of full gene deletions remained comparable at 5%. Comparison with pb-StarPhase revealed discrepancies in 12 of 94 assemblies in the release 1 dataset. For instance, in sample HG02257, Aldy, Cyrius, and deCYPher consistently identified the genotype as *2/*35, whereas pb-StarPhase reported *2/*2. Notably, the *35-defining variants were present in the BAM and VCF files in the pb-StarPhase pipeline, but the local read depth over the *35-specific region was only 5x in HG02257-p, suggesting that the misclassification likely resulted from insufficient coverage - a known limitation of pb-StarPhase under low-depth conditions.

## Introduction

Pharmacogenomics investigates the genetic factors influencing drug efficacy and toxicity, with pharmacogenes identified as key contributors to therapeutic outcomes. The Pharmacogene Variation (PharmVar) Consortium curates star-allele definitions and structural-variant annotations to enable reproducible allele calling (Gaedigk et al., 2018), while CPIC and PharmGKB provide evidence-based drug-dosing and reporting guidance that depends on accurate genotype calls (Relling & Klein, 2011; Whirl-Carrillo et al., 2021). PharmVar currently catalogues pharmacogenes such as *CYP1A2, CYP2A6, CYP2A13, CYP2B6, CYP2C8, CYP2C9, CYP2C19, CYP2D6, CYP3A4, CYP3A5, CYP4F2, DPYD, NUDT15, NAT2*, and *SLCO1B1*, offering standardized allele definitions across major drug-metabolizing enzymes and transporters. Concurrently, CPIC has published clinical guidelines for an expanded array of genes—including *ABCG2, CACNA1S, CFTR, CYP2B6, CYP2C8, CYP2C9, CYP2C19, CYP2D6, CYP3A4, CYP3A5, CYP4F2, DPYD, G6PD, HLA-A, HLA-B, HLA-C, IFNL3, IFNL4, MT-RNR1, NUDT15, RYR1, SLCO1B1, TPMT, UGT1A1*, and *VKORC1*. Collectively, these complementary resources establish a molecular basis for the integration of pharmacogenomics into clinical practice.

This tool enables comprehensive analysis of all PharmVar level 1A genes. Within this group, cytochrome P450 (CYP) enzymes play a significant role, as they are involved in phase I oxidative metabolism for many commonly prescribed drugs and contribute to interindividual differences in pharmacokinetics and drug response (Zanger & Schwab, 2013). Reviews of the CYP superfamily summarize how genetic variation and regulation of expression in these enzymes shape drug metabolism phenotypes and influence efficacy, adverse drug reactions, and drug–drug interactions.

Among CYP and related pharmacogenes, loci such as *CYP2D6, CYP2B6, CYP2C9, CYP2C19*, and others are highly polymorphic. They are well-established predictors of clinically actionable metabolic phenotypes (for example, poor, intermediate, normal, rapid, or ultrarapid metabolizers) used in therapeutic decision-making. These genotype–phenotype translations underpin numerous Clinical Pharmacogenetics Implementation Consortium (CPIC) guidelines and PharmGKB clinical annotations that convert molecular test results into prescribing recommendations.

Accurate star-allele assignment for these genes is complicated by multiple genetic variation types. In addition to single-nucleotide variants (SNVs) and small indels, clinically important structural variation occurs frequently — including whole-gene deletions and duplications, hybrid alleles created by gene–pseudogene conversions, and complex rearrangements — all of which can have large effects on enzyme activity. *CYP2D6* serves as a classic example; it is located in a region with related pseudogenes such as *CYP2D7* and features numerous structural alterations. Approximately 5-10% of people have copy-number variants like gene deletions or duplications, while *CYP2D6/CYP2D7* hybrid alleles are found in about 2-6%. Altogether, these variations make up more than 10% of clinically significant differences in *CYP2D6* (Hicks et al., 2015; Relling & Klein, 2011; Whirl-Carrillo et al., 2021).

Conventional short-read sequencing and basic variant-calling pipelines frequently misidentify structural variants, leading to inaccurate star-allele calls and incorrect downstream enzyme activity assignments. Consequently, substantial inter-laboratory and inter-algorithm discrepancies in phenotype assignment often originate from differences in the underlying genotype calls, particularly when structural variation is present (Del Tredici et al., 2018; Gaedigk et al., 2018). A host of computational tools and experimental workflows have been developed to tackle pharmacogene genotyping. Short-read–based star-allele callers have achieved high concordance for common alleles (Chen et al., 2021; Hari et al., 2023; Lee et al., 2019; Twesigomwe et al., 2020), but their effectiveness is limited when complex structural rearrangements or high sequence homology to pseudogenes are involved. Recent benchmarking studies indicate that sequence ensemble approaches combined with specialized algorithms perform well. However, complete concordance among all allele classes—including rare structural variants and hybrids—has not been consistently attained (Chen et al., 2021; Qiao et al., 2016). Integrating multiple sequencing techniques, including PacBio HiFi, ONT ultra-long reads, and Illumina Hi-C reads, to construct haplotype-resolved assemblies has demonstrated the ability to resolve complex loci, detect novel structural variants, and provide contiguous haplotypes that are essential for unambiguous allele definition (Cheng et al., 2021; Liao et al., 2023; Nurk et al., 2022).

The pb-StarPhase demonstrates a phase-aware pipeline diploid typing of twenty-one clinically important pharmacogenes utilizing PacBio HiFi reads, with the ability to correct or refine historical genotype calls (Holt et al., 2024). Although pb-StarPhase have advanced allele resolution for complex pharmacogenes, their performance can still be affected by sequencing coverage and read length, which may lead to incomplete or uncertain genotype calls under suboptimal data quality. These limitations highlight the need for a targeted and systematic framework that operates consistently across clinically relevant pharmacogenes, integrates PharmVar allele definitions with CPIC and PharmGKB conventions, and delivers high-confidence SNV, CNV, and hybrid allele calls directly from haplotype-resolved genome assemblies.

We therefore developed **deCYPher**, a computational method for star-allelic resolution of pharmacogenes listed in CPIC/PharmGKB/PharmVar using haplotype-resolved long-read assemblies, particularly in genes with complex structural variations. We benchmarked deCYPher on nine PharmVar 1A-level genes (*CYP2D6, CYP2B6, CYP2C9, CYP2C19, CYP3A5, CYP4F2, DPYD, NUDT15*, and *SLCO1B1*) and compared its results with those of Aldy, Cyrius, pb-StarPhase, and TypeAssembly. Compared to existing short-read or long-read mapping–based approaches, deCYPher leverages fully assembled haplotypes to achieve high accuracy and unambiguous allele definitions, thereby enhancing the reliability of clinical pharmacogenomic reporting.

## Results

### Method Overview

The 47 phased diploid genome assemblies from the Human Pangenome Reference Consortium (HPRC), were analyzed with our **deCYPher** pipeline (Fig.1). The deCYPher was designed to achieve the highest resolution for allele assignments defined by the PharmVar Consortium. The standardized CYP nomenclature — comprising the *CYP* root, family number, subfamily letter, gene number, and a star allele designation — was used to ensure consistency with established pharmacogenomic conventions.

**Figure 1.**
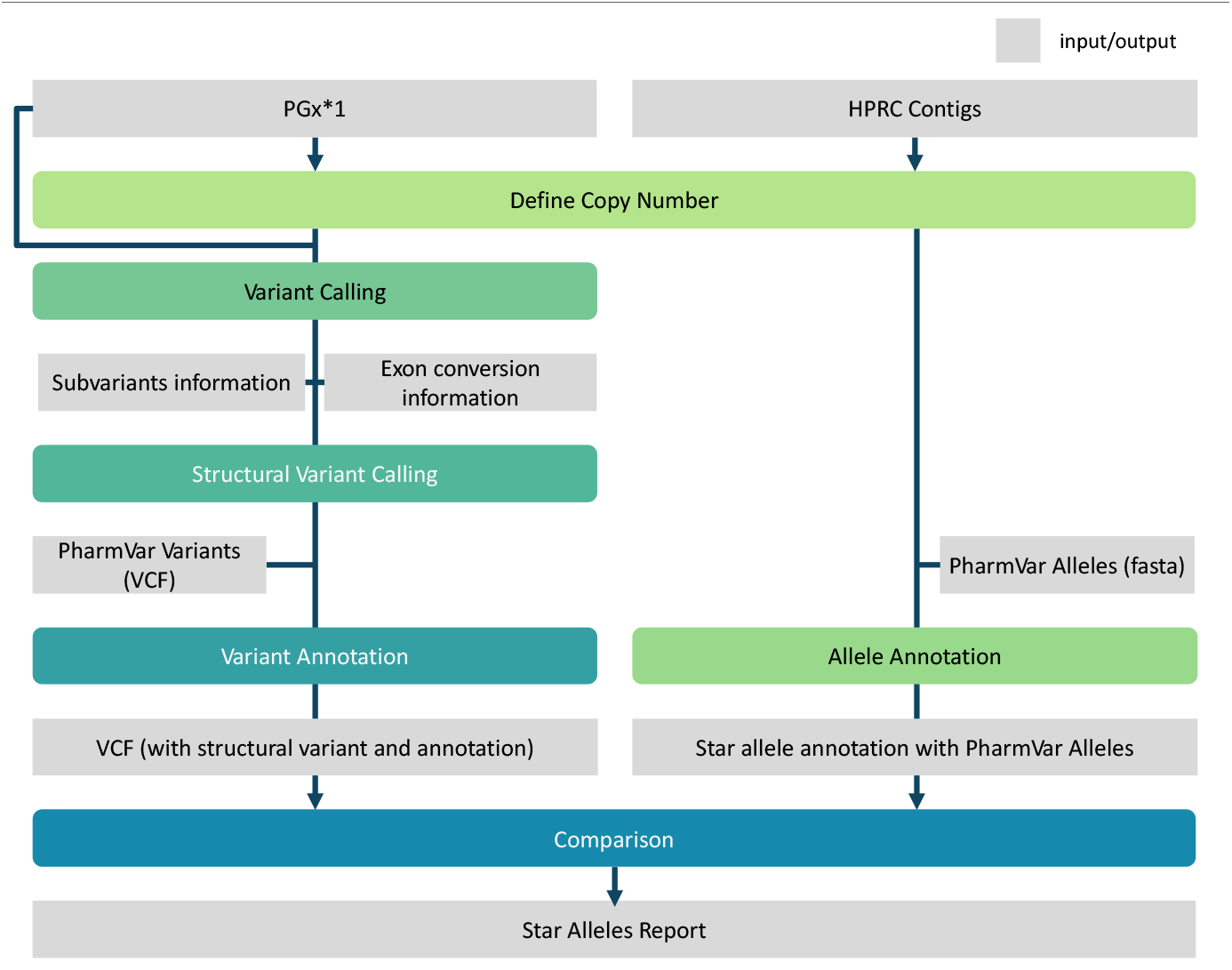
Workflow of the deCYPher pipeline for CYP allele annotation in diploid genome assemblies. The pipeline integrates the long-read alignment tool minimap2 (Li, 2018) with custom Python and Shell scripts to map allele sequences from the Pharmacogene Variation Consortium (PharmVar) database onto genome assemblies, including the HPRC Year-1 and Year-2 releases. This approach enables accurate identification of CYP gene alleles on phased contigs, achieving up to 95% accuracy in high-resolution allele assignment.

For each haplotype-resolved assembly, deCYPher first identified the intervals of target pharmacogenes and mapped curated reference sequences representing star alleles (“star-one” references) to the phased contigs. Alignments were performed at full-sequence resolution to capture both single-nucleotide variants (SNVs) and small indels. Alleles matching known PharmVar definitions were directly assigned. When variant patterns did not match any existing alleles, deCYPher flagged them as potential novel alleles, with particular attention to previously unreported structural variants (SVs) such as novel hybrid genes or deletion genes.

We validated deCYPher-identified structural variants by comparing them to genotypes from Aldy, Cyrius, and pb-StarPhase. Cross-comparison confirmed the structural variants.

### Overview of the HPRC CYP Year-1 and Year-2 releases haplotypes and phenotypes

We fully characterized the high-resolution alleles of CYP genes for all known CYP alleles in Year-1 47 HPRC human genome assemblies and Year-2 232 HPRC human genome assemblies (Fig.2A-B). According to the CYP genotype definitions from the Pharmacogene Variation Consortium (PharmVar; https://www.pharmvar.org/), the CYP2D6 gene includes 185star alleles, encompassing numerous hybrid and deletion alleles, as well as copy number variants (CNVs) (Taylor et al., 2020). Our results show that, among the 94 CYP2D6 haplotypes in the HPRC Year-1 dataset, 6 haplotypes (6.38%, 6/94) harbor one or more CNVs. Additionally, 6 haplotypes (6.38%, 6/94) were identified as full gene deletions, as illustrated in Figure 2C. Similarly, in the HPRC Year-2 dataset, 66 out of 464 CYP2D6 haplotypes (14.2%, 66/464) contained one or more CNVs, while 22 haplotypes (4.74%, 22/464) were classified as full gene deletions, as shown in Figure 2C.

**Figure 2.**
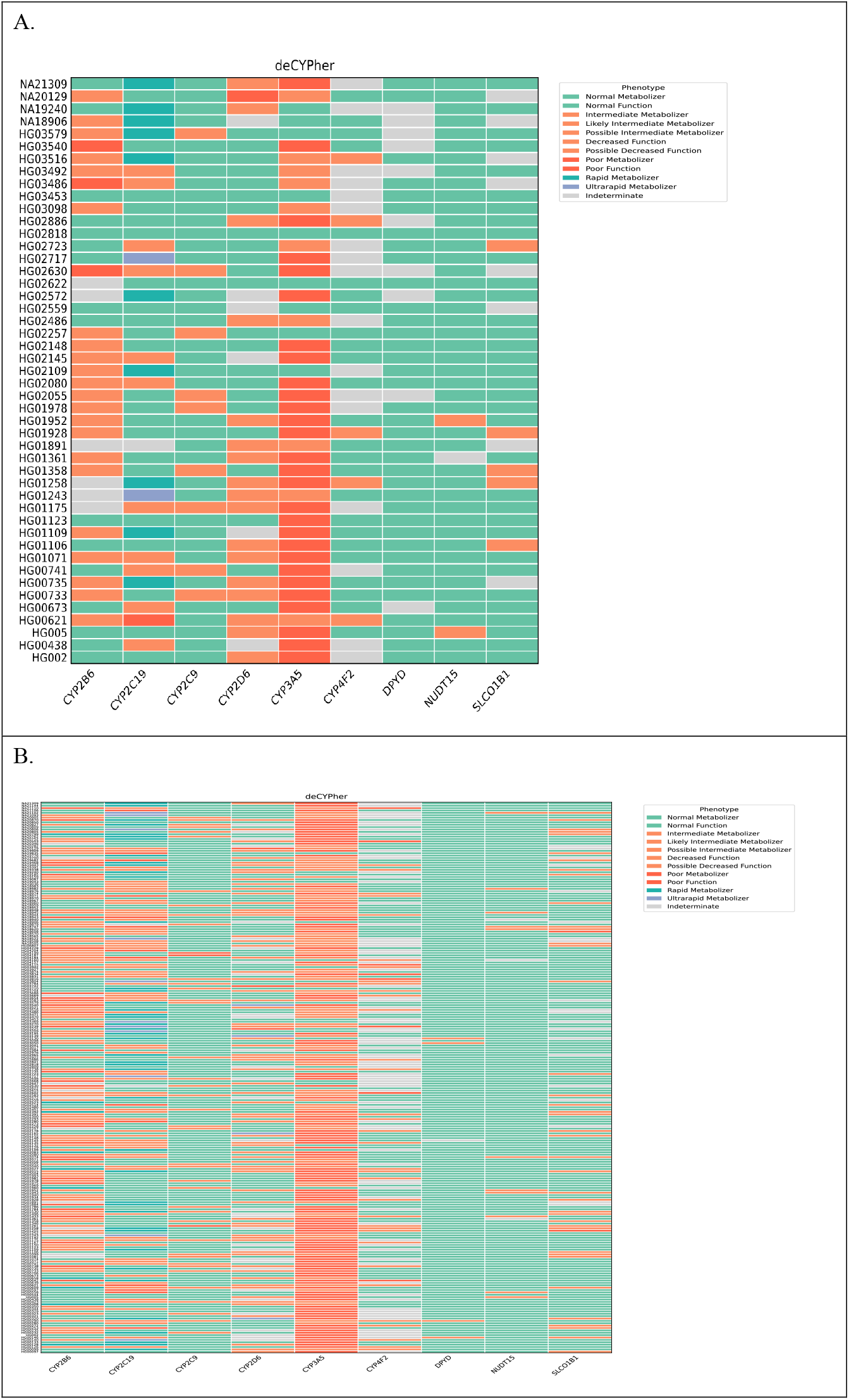
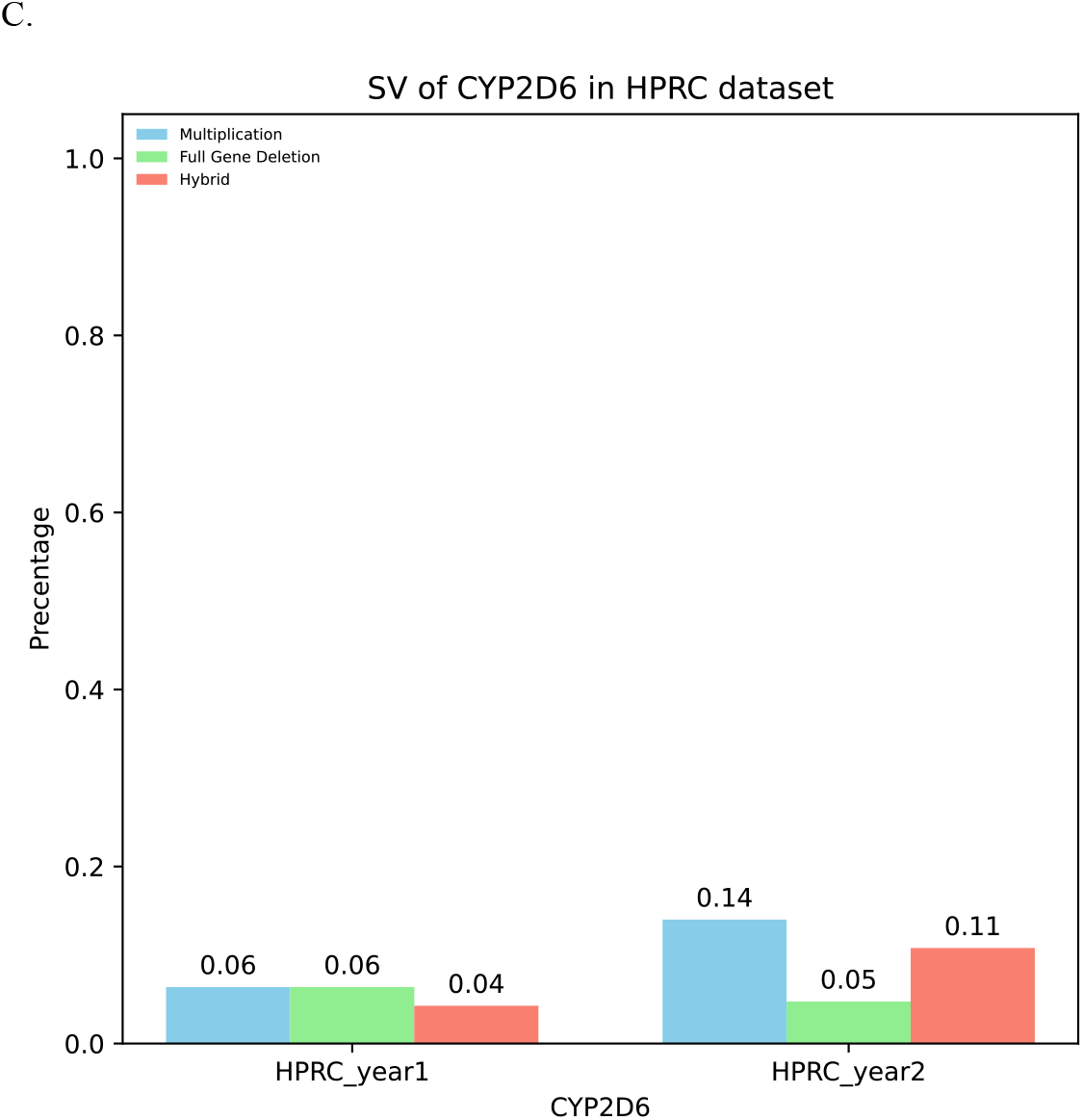
Phenotypic profiles of pharmacogenes and structural variants (SVs) of *CYP2D6* identified in the 47 phased assemblies and 232 phased assemblies analyzed by deCYPher. (A)Heatmap of predicted pharmacogene phenotypes across the HPRC Year-1 cohort. (B)Heatmap of predicted pharmacogene phenotypes across the HPRC Year-2 cohort. (C)Structural variant configurations of *CYP2D6*, including full-gene deletions, duplications, and gene–pseudogene hybrid arrangements, as detected in the HPRC Year-1,-2 assemblies.

Regarding phenotype predictions, 24 samples (51.06%, 24/47) in the HPRC Year-1 assemblies were classified as Normal Metabolizers, whereas in the HPRC Year-2 assemblies, 121 samples (52.16%, 121/232) fell into this category (detailed data are provided in Supplemental Tables 1 and 2). These findings address the limitations of previous technologies, which could not resolve CYP gene structural variations at base-pair resolution across whole-genome sequences, highlighting the importance of deCYPher for the comprehensive characterization of pharmacogene genetic variation.

In this study, we denote paternal haplotypes as SampleID-p and maternal haplotypes as SampleID-m, where SampleID corresponds to the individual sample. For example, HG002-p and HG002-m represent the paternal and maternal haplotypes of sample HG002, respectively.

Within the HPRC assemblies, in addition to *CYP2D6*, we also identified structural variants in other pharmacogenes. For example, in HG01993-m we detected a *CYP2B6* hybrid gene, while in HG00423-m we observed a full-gene deletion of *CYP3A5* — a variant that has never been previously reported. Furthermore, in HG02615-m we identified a CYP4F2 copy number variation (CNV), which is also novel, and in HG02615-p we found a full-gene deletion of CYP4F2. Notably, all of these non-*CYP2D6* structural variants were discovered in the HPRC Year-2 phased assemblies. This suggests that expanding the size of cohorts with PacBio HiFi phased assemblies may enable the discovery of additional low allele frequency and novel structural variants, further underscoring the importance of **deCYPher** for structural variant analysis in pharmacogenes (Supplemental Tables 3 and 4).

### Distinctive variant positions defining the CYP2D6*68 hybrid allele

We identified a *CYP2D6* hybrid allele, CYP2D6 *68, in five phased assemblies from the HPRC dataset: HG00733-p, HG01258-p, HG01784-m, NA20805-p, and NA20827-m.

According to PharmVar, *68 is defined by a CYP2D7-to-CYP2D6 gene conversion spanning exons 2 to 9. By contrast, *\**36 is characterized by a conversion limited to exon 9, and *82 by a conversion restricted to exon 2. Consequently, any haplotype carrying the exon 2–9 conversion of *68 will inherently meet the defining criteria of both *36 (exon 9 conversion) and *82 (exon 2 conversion). Based on this relationship, our initial strategy was to classify any assembly carrying both *36 and *82 as *68 (Fig. 3A). Using this approach, we detected three *68 alleles in the HPRC Year-1 phased assemblies—one in HG00733-p and two in HG01258-p—yielding results that were fully concordant with calls from Aldy and Cyrius.

**Figure 3.**
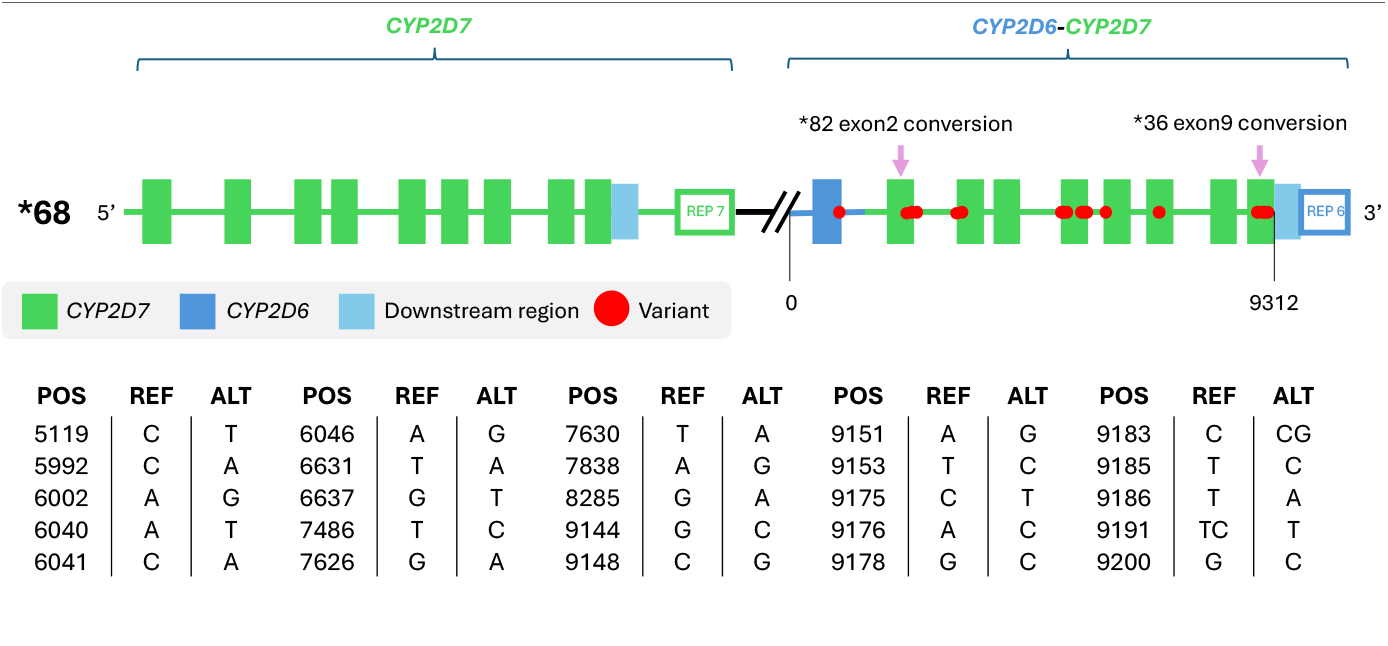
Detection and variant profile of CYP2D6*68. CYP2D6*68 is characterized by a continuous CYP2D7-to-CYP2D6 conversion spanning exons 2–9, in contrast to *36 (exon 9 only) and *82 (exon 2 only). Assemblies harboring both *36 and *82 were classified as *68. Variant positions defining the *68 hybrid allele across exons 2–9 are shown, with each marker indicating a characteristic nucleotide change.

Although carrying both *36 and *82 as a *68 criterion successfully identified *68, it may not capture all instances of this hybrid allele, as *68 involves a continuous exon 2–9 conversion. To refine detection, we sought variant positions that could serve as a more complete signature for *68. We first compiled all star alleles present across confirmed *68 assemblies and found consistent co-occurrence of the following alleles: *10, *36, *39, *49, *53, *74, *82, *83, *86, *88 (with one variant site optionally absent), *131, *132, *139, and *176. We then recorded the variant positions corresponding to these alleles, excluding NG_008376.4:6032 T>C, to generate a representative *68 variant profile (VCF) (Fig. 3B and Supplemental Table 5).

### Fusion of *CYP2B6*

Previous studies have shown that the **CYP2B6*29** allele shares complete sequence identity with exons 1–4 of ***CYP2B7*** and exons 5–9 of ***CYP2B6***, and is associated with reduced enzymatic activity. In the PharmVar database, CYP2B6*29 is annotated as a hybrid gene between *CYP2B7* and *CYP2B6*. Furthermore, the GenBank database contains FASTA sequences representing the *CYP2B7/CYP2B6* breakpoint junction, as documented in earlier reports (Rotger *et al*. 2007; Martis *et al*. 2013; Gaedigk *et al*. 2018).

Although the genomic characterization of **CYP2B6*29** is already well documented, current genotyping tools such as **Cyrius, Aldy**, and **pb-StarPhase** perform suboptimally in detecting this allele. In both the HPRC Year-1 and Year-2 phased assemblies, no samples were classified as CYP2B6*29 by these tools. However, when applying **deCYPher** to the HPRC Year-2 phased assemblies, we identified **HG01993-m** as carrying the CYP2B6*29 allele.

deCYPher recognized this allele based on two key observations. First, we detected abnormal mapping patterns when aligning the HG01993-m contigs to the **CYP2B6*1** reference: read mapping with generally poor mapping quality, except for the primary mapping segment spanning **exons 5–9**, where mapping quality was higher, whereas **exons 1–4** were predominantly soft-clipped (Fig. 4A). This mapping profile is consistent with the known structure of CYP2B6*29. Second, when aligning HG01993-m to the *CYP2B7/CYP2B6* breakpoint junction sequence, we observed almost perfect alignment across the junction with no evidence of soft or hard clipping, and only a few single-nucleotide variants (SNVs) (Fig. 4B). These Integrative Genomics Viewer (IGV)-based visualizations (Robinson et al., 2011) provide orthogonal evidence that deCYPher correctly identified HG01993-m as CYP2B6*29. Existing algorithms missed this call. This demonstrates deCYPher’s unique ability to resolve hybrid alleles in pharmacogenes that are often misclassified or overlooked by short-read–based tools.

**Figure 4.**
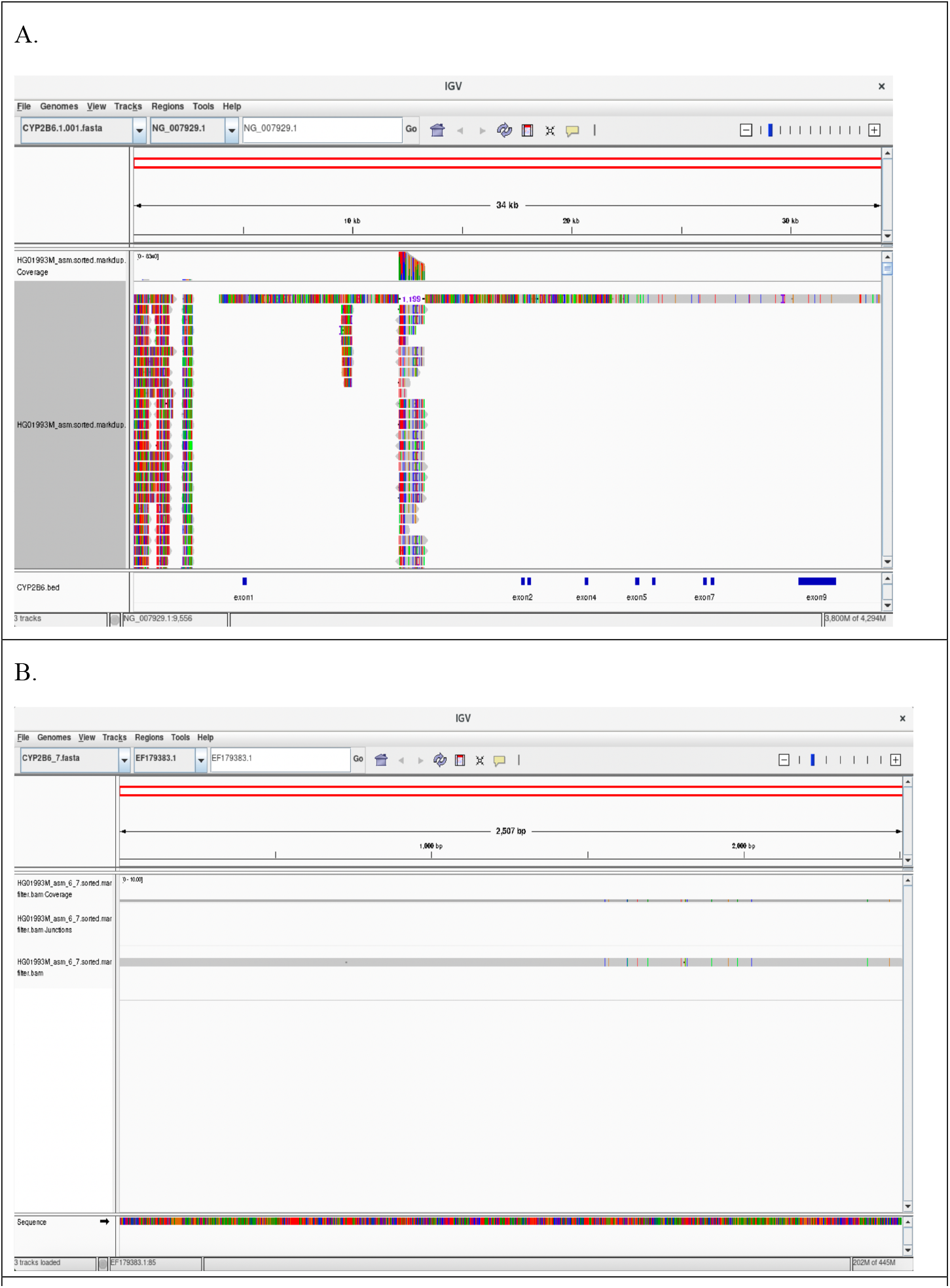
IGV visualization of CYP2B6*29 in HG01993-m. (A) IGV alignment view of the HG01993-m contigs against the CYP2B6*1 reference. The region corresponding to exons 1–4 displayed extensive soft clipping and poor mapping quality, whereas alignment improved across exons 5–9, consistent with the CYP2B6*29 hybrid structure.

### Comparisons with pb-StarPhase in HPRC genomes

pb-StarPhase (Holt et al., 2024) provides an innovative approach for pharmacogene genotyping using PacBio long-read whole-genome sequencing. To directly compare its performance with deCYPher, we applied the recommended pb-StarPhase pipeline to the 47 HPRC Year-1 samples, starting from randomly selected long-read raw data. We then generated diplotype calls for all samples and compared them against deCYPher results.

During this comparison, we observed that pb-StarPhase frequently produced “NO_MATCH” calls. For the purpose of evaluation, such calls were treated as discordances and provisionally considered errors on the pb-StarPhase side. For other discordant cases, we examined upstream evidence (i.e., BAM and VCF files) to adjudicate correctness. Specifically, if a critical variant was clearly present in the alignments and variant calls but missing from the output, we classified the call as incorrect. Conversely, if a critical variant was clearly present in the alignments and variant calls and also present in the output, we classified the call as correct. In cases where both tools produced plausible but differing outputs supported by partial evidence, we labeled the discrepancy as “Unknown”, reflecting unresolved classification.

As an illustrative example, in HG03516, both the pbmm2-aligned bam file and the Deepvariant vcf file contained the variant chr22:42129770 G>A, a defining variant of CYP2D6*17 (Fig. 5). Nevertheless, pb-StarPhase assigned this sample as CYP2D6*2/*2, missing the variant, whereas deCYPher correctly identified the diplotype as CYP2D6*17/*2.

**Figure 5.**
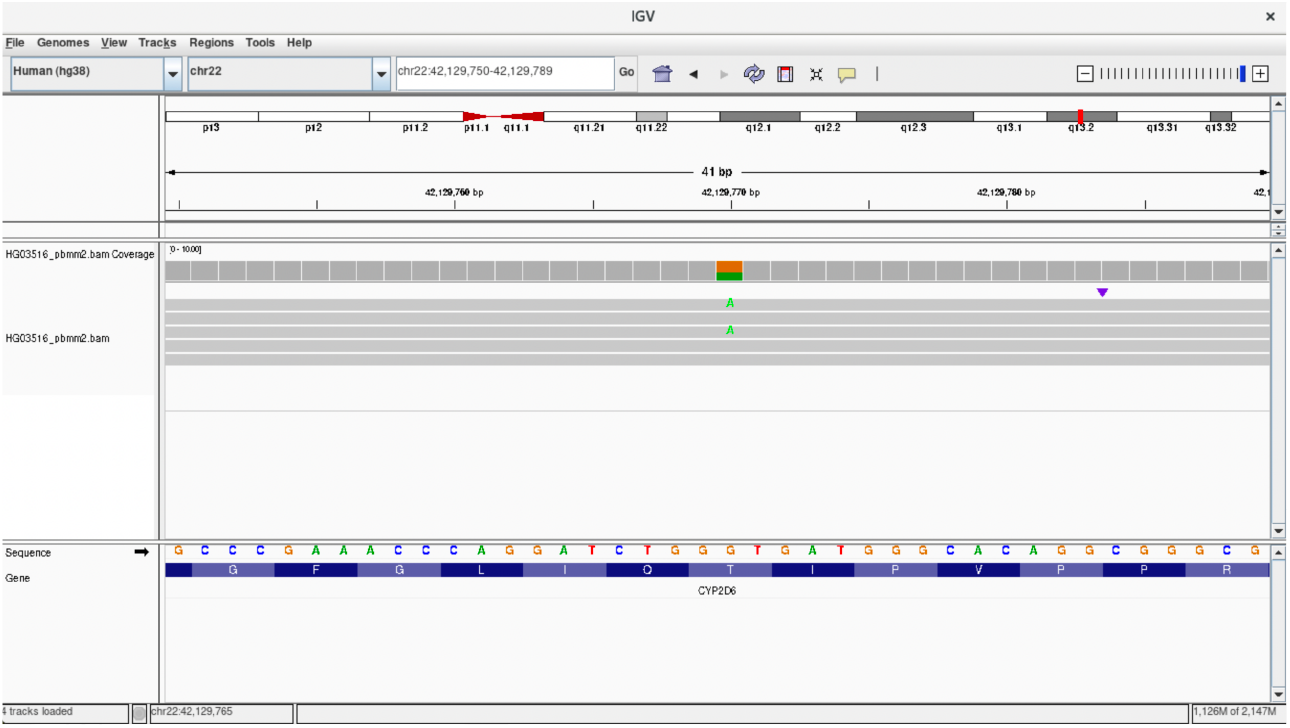
IGV visualization of the CYP2D*17-defining variant in HG03516. This track shows clear evidence of the chr22:42129770 G>A substitution. Despite this, pb-StarPhase assigned the sample as CYP2D6*2/*2, whereas deCYPher correctly identified it as CYP2D6*17/*2.

Overall, deCYPher demonstrated superior accuracy for *CYP2D6*, achieving a precision of 94.68%, compared to 87.23% for pb-StarPhase — the largest discrepancy observed among the nine pharmacogenes analyzed. Notably, both tools produced identical results for *CYP2C9* and *NUDT15* (Table 1).

**Table 1.**
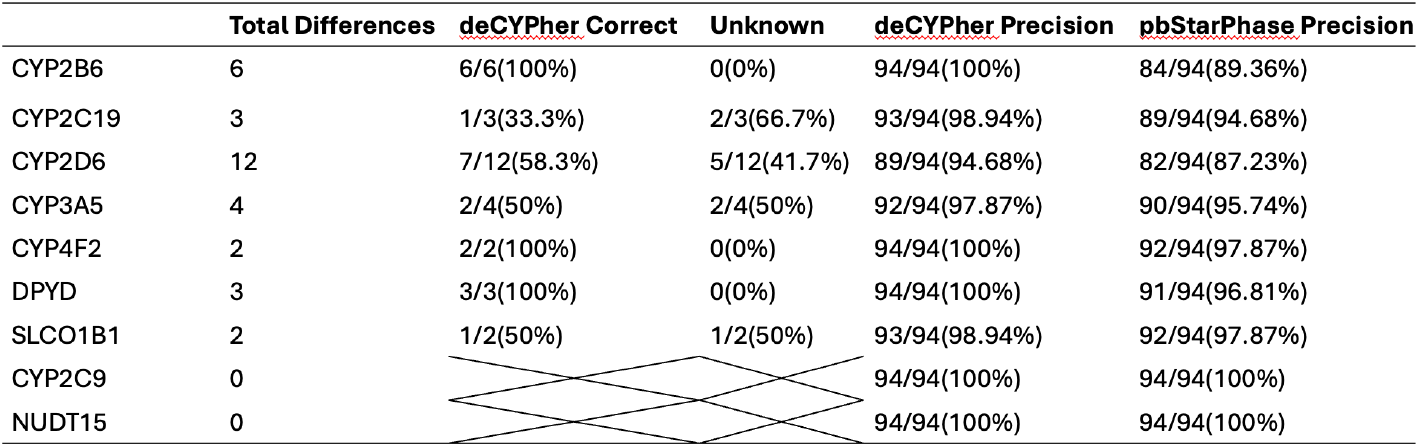
Comparison between deCYPher and pb-StarPhase.

These findings highlight the limitations of pb-StarPhase when allele-defining, while underscoring the advantage of deCYPher in leveraging haplotype-resolved assemblies to deliver unambiguous star-allele assignments. This accuracy is particularly critical for complex loci such as *CYP2D6*, where reliable genotype-to-phenotype translation is essential for clinical pharmacogenomics.

## Discussion

In this study, we developed and applied **deCYPher**, a computational framework for star allele typing of pharmacogenes using haplotype-resolved assemblies. By leveraging long-read contiguity and phasing, deCYPher enables accurate assignment of PharmVar-defined alleles, detection of structural variants, and standardized diplotype-to-phenotype translation consistent with CPIC conventions. Compared to existing genotyping tools such as Aldy, Cyrius, and pb-StarPhase (Chen et al., 2021; Hari et al., 2023; Holt et al., 2024), deCYPher demonstrated improved accuracy, particularly in loci with complex structural variation.

Hybrid alleles of *CYP2D6*, such as *36 and *82, have been well characterized in previous studies (Gaedigk et al., 2018; Gaedigk et al., 2017). In line with these reports, we identified CYP2D6*68, a complex hybrid allele defined in PharmVar as a gene conversion event spanning exons 2–9 from *CYP2D7* (Gaedigk et al., 2018). Importantly, our analysis also revealed structural variants outside of *CYP2D6*, including a CYP2B6*29 hybrid in HG01993-m, which, to our knowledge, has not been reported in other tools before. These findings highlight the strength of deCYPher in resolving both known and unclear structural variants across pharmacogenes, particularly when complex scenarios involving CNVs, hybrid genes, or large deletions complicate haplotype assignment.

These findings highlight a broader limitation of short-read pipelines. Previous work has shown that genotype-to-phenotype assignments can differ substantially across laboratories and algorithms, especially in the presence of structural variants (Twesigomwe et al., 2020). In such cases, ambiguity often arises not because phenotype rules differ, but because the underlying genotype calls are inconsistent. By integrating haplotype-resolved assemblies with PharmVar allele definitions, deCYPher substantially reduces this discordance and produces more reliable haplotype calls.

Our benchmarking against HPRC assemblies further supports this advantage. While pb-StarPhase achieved a precision of 8% for *CYP2D6* calls, deCYPher achieved 94.7%, representing the largest performance gap across the nine genes evaluated. In two genes (*CYP2C9* and *NUDT15*), both tools produced identical results, indicating that the primary benefit of deCYPher emerges in genes with extensive structural complexity. These results are consistent with prior observations that structural variation is a major source of error in pharmacogene genotyping (Twesigomwe et al., 2020; Zhou et al., 2017).

Finally, our work parallels advances in other structurally complex genomic loci, such as the KIR region. The SKIRT pipeline has demonstrated how leveraging high-quality assemblies enables the discovery of novel gene fusions and CNVs in KIR genes (Hung et al., 2024; Zhou et al., 2024). By analogy, deCYPher provides a foundation for high-resolution pharmacogene genotyping, paving the way for future studies in larger HiFi-based cohorts and the systematic discovery of rare or novel pharmacogene SVs.

## Methods

### deCYPher - Algorithms of the Pharmacogenes annotation Pipeline

#### Read Mapping and Variant Detection

The deCYPher pipeline was developed to resolve PharmVar-defined star alleles and structural variation, including gene duplications and deletions. To capture multiple polymorphic alleles at each locus, we aligned PharmVar reference allele sequences to haplotype-resolved genome assemblies using **minimap2** (Li, 2018) optimized for PacBio HiFi data. Parameter settings and alignment configurations are provided in **Supplementary Note 2**. To validate structural variants identified by deCYPher, we compared the results with genotypes obtained from established tools, including Aldy, Cyrius, and pb-StarPhase. The input data consisted of short-read Illumina whole-genome sequencing (WGS) from the 1KGP and HPRC cohorts, as well as long-read PacBio HiFi sequencing. After cross-comparing these results, we evaluated and confirmed the presence of structural variants.

#### Output and Visualization

deCYPher reports haplotype-level star allele calls in a structured JSON format, providing a standardized and machine-readable representation of nine PharmVar/CPIC-prioritized pharmacogenes. This format encodes PharmVar-defined star alleles together with copy number information and annotations of putative novel structural variants, thereby supporting downstream analysis, integration into pharmacogenomic pipelines, and clinical interpretation. Further details regarding the output structure and visualization are provided in **Supplementary Note 1**.

## Data access

All data generated or analyzed during this study are included in this article, its Supplemental files, and Zenodo (https://doi.org/10.5281/zenodo.8094803, https://zenodo.org/records/14854401). deCYPher source code is available on GitHub (https://github.com/HolidayChang/deCYPher).

## Competing interest statement

The authors declare no competing interests.

### Acknowledgment

This work was supported in part by computational and storage resources provided by the National Center for High-performance Computing (NCHC), National Applied Research Laboratories (NARLabs), Taiwan. Funding informaiton

## Author contributions

J.S.H., P-L.C., Y-S.L., and T-K.H. proposed the initial idea of annotating pharmacogene alleles using high-quality assemblies, initiated the project, and provided input during discussions.

T-Y.C. conceived the study, designed the analysis pipeline, acquired the HPRC data, performed the analyses, interpreted the results, and generated the figures and tables.

T-Y.C. also managed the writing and organization of the manuscript, with suggestions and proofreading from J.S.H.

J.S.H. supervised the overall project.

All authors read and approved the final manuscript.

## Supplemental material

**Supplemental File1:** decepher.xlsx

**Supplemental Table 1.** Predicted phenotypes for pharmacogenes in the HPRC Year-1 phased assemblies as determined by **deCYPher**.

| HPRC Year1 | Ultrarapid Metabolizer, Rapid Metabolizer | Normal Metabolizer, Normal Function | Intermediate Metabolizer, Likely Intermediate Metabolizer, Possible Intermediate Metabolizer, Decreased Function, Possible Decreased Function | Poor Metabolizer, Poor Function | Indeterminate |
| --- | --- | --- | --- | --- | --- |
| CYP2B6 | 0/47(0%) | 15/47(31.91%) | 24/47(51.06%) | 3/47(6.38%) | 5/47(10.64%) |
| CYP2C9 | 0/47(0%) | 38/47(33.3%) | 12/47(66.7%) | 0/47(0%) | 0/47(0%) |
| CYP2C19 | 14/47(29.79%) | 20/47(42.55%) | 12/47(25.53%) | 1/47(2.13%) | 0/47(0%) |
| CYP2D6 | 1/47(2.13%) | 24/47(51.06%) | 17/47(36.17%) | 1/47(2.13%) | 4/47(8.51%) |
| CYP3A5 | 0/47(0%) | 6/47(12.77%) | 11/47(0%) | 30/47(100%) | 0/47(0%) |
| CYP4F2 | 0/47(0%) | 25/47(53.19%) | 5/47(10.64%) | 0/47(0%) | 17/47(36.17%) |
| DPYD | 0/47(93.62%) | 44/47(93.62%) | 0/47(0%) | 0/47(0%) | 3/47(6.38%) |
| SLCO1B1 | 0/47(0%) | 35/47(74.47%) | 5/47(10.64%) | 0/47(0%) | 7/47(14.89%) |
| NUDT15 | 0/47(0%) | 44/47(93.62%) | 2/47(4.26%) | 0/47(0%) | 1/47(2.13%) |

**Supplemental Table 2.** Predicted phenotypes for pharmacogenes in the HPRC Year-2 phased assemblies as determined by **deCYPher**.

| HPRC Year2 | Ultrarapid Metabolizer, Rapid Metabolizer | Normal Metabolizer, Normal Function | Intermediate Metabolizer, Likely Intermediate Metabolizer, Possible Intermediate Metabolizer, Decreased Function, Possible Decreased Function | Poor Metabolizer, Poor Function | Indeterminate |
| --- | --- | --- | --- | --- | --- |
| CYP2B6 | 10/232(4.31%) | 83/232(35.78%) | 96/232(41.38%) | 31/232(13.36%) | 12/232(5.17%) |
| CYP2C9 | 0/232(0%) | 189/232(81.47%) | 40/232(17.24%) | 3/232(1.29%) | 0/232(0%) |
| CYP2C19 | 54/232(23.28%) | 89/232(38.36%) | 73/232(31.47%) | 14/232(6.03%) | 2/232(0.86%) |
| CYP2D6 | 4/232(1.72%) | 121/232(52.16%) | 72/232(31.03%) | 3/232(1.29%) | 32/232(13.79%) |
| CYP3A5 | 0/232(0%) | 27/232(11.64%) | 87/232(37.5%) | 118/232(50.86%) | 0/232(0%) |
| CYP4F2 | 0/232(0%) | 108/232(46.55%) | 34/232(14.66%) | 8/232(3.45%) | 82/232(35.34%) |
| DPYD | 0/232(0%) | 227/232(97.84%) | 4/232(1.72%) | 0/232(0%) | 1/232(0.43%) |
| SLCO1B1 | 0/232(0%) | 170/232(73.28%) | 40/232(17.24%) | 2/232(0.86%) | 20/232(8.62%) |
| NUDT15 | 0/232(0%) | 218/232(93.97%) | 11/232(4.74%) | 0/232(0%) | 3/232(1.29%) |

**Supplemental Table 3,4.** Predicted structural variants for pharmacogenes in the HPRC Year-1 and Year-2 phased assemblies as determined by **deCYPher**.

| HPRC Year1 | CYP2B6 | CYP2C9 | CYP2C19 | CYP2D6 | CYP3A5 | CYP4F2 | DPYD | SLCO1B1 | NUDT15 |
| --- | --- | --- | --- | --- | --- | --- | --- | --- | --- |
| CNV | 0/94(0%) | 0/94(0%) | 0/94(0%) | 6/94(6.38%) | 0/94(0%) | 0/94(0%) | 0/94(0%) | 0/94(0%) | 0/94(0%) |
| Full Gene Deletion | 0/94(0%) | 0/94(0%) | 0/94(0%) | 6/94(6.38%) | 0/94(0%) | 0/94(0%) | 0/94(0%) | 0/94(0%) | 0/94(0%) |
| Hybrid | 0/94(0%) | 0/94(0%) | 0/94(0%) | 4/94(4.26%) | 0/94(0%) | 0/94(0%) | 0/94(0%) | 0/94(0%) | 0/94(0%) |
| SNV | 94/94(100%) | 94/94(100%) | 94/94(100%) | 82/94(87.23%) | 94/94(100%) | 94/94(100%) | 94/94(100%) | 94/94(100%) | 94/94(100%) |

| HPRC Year2 | CYP2B6 | CYP2C9 | CYP2C19 | CYP2D6 | CYP3A5 | CYP4F2 | DPYD | SLCO1B1 | NUDT15 |
| --- | --- | --- | --- | --- | --- | --- | --- | --- | --- |
| CNV | 0/464(0%) | 0/464(0%) | 0/464(0%) | 65/464(14%) | 0/464(0%) | 1/464(0.22%) | 0/464(0%) | 0/464(0%) | 0/464(0%) |
| Full Gene Deletion | 0/464(0%) | 0/464(0%) | 0/464(0%) | 22/464(4.74%) | 1/464(0.22%) | 1/464(0.22%) | 0/464(0%) | 0/464(0%) | 0/464(0%) |
| Hybrid | 1/464(0.22%) | 0/464(0%) | 0/464(0%) | 50/464(10.78%) | 0/464(0%) | 0/464(0%) | 0/464(0%) | 0/464(0%) | 0/464(0%) |
| SNV | 463/464(99.78%) | 464/464(100%) | 464/464(100%) | 375/464(80.82%) | 463/464(99.78%) | 462/464(99.57%) | 464/464(100%) | 464/464(100%) | 464/464(100%) |

